# Trait mindfulness predicts inter-brain coupling during naturalistic face-to-face interactions

**DOI:** 10.1101/2021.06.28.448432

**Authors:** Phoebe Chen, Ulrich Kirk, Suzanne Dikker

## Abstract

In recent years, the benefits of practicing mindfulness have raised much public and academic interest. Mindfulness emphasizes cultivating awareness of our immediate experience, and has been associated with compassion, empathy and various other prosocial traits. However, experimental evidence pertaining to its prosocial benefits in social settings is lacking. In this study, we investigate neural correlates of trait mindfulness during naturalistic dyadic interactions, using both individual brain and inter-brain coupling measures. We used the Muse headset, a portable electroencephalogram (EEG) device, to record participants’ brain activity during a ∼10 minutes’ naturalistic dyadic interaction (N = 62) in an interactive art setting. They further completed the Mindful Attention Awareness Scale (MAAS) and the Interpersonal Reactivity Index (IRI). This allowed us to ask whether inter-brain coupling during naturalistic interactions can be predicted by dyads’ affective traits and trait mindfulness, respectively. First, we failed replicating prior laboratory-based findings with respect to individual brain responses as they relate to mindfulness. Trait mindfulness *did, however*, predict inter-brain coupling within dyads, in theta (∼5-8 Hz, p < 0.001) and beta frequencies (∼26-27Hz, p < 0.001). Finally, we found a negative correlation between personal distress and trait mindfulness (t(475) = -5.493, p < 0.001). These findings underscore the importance of conducting social neuroscience research in ecological settings and enrich our understanding of multi-brain neural correlates of mindfulness during social interaction, while raising critical practical considerations regarding the viability of commercially available EEG systems.

## Introduction

Recent years have seen an increase in popular interest in the benefits of practicing mindfulness. As a personality trait, mindfulness refers to attending to the present moment experience without judging occurring feelings or thoughts (Bishop et al., 2006), and has been associated with prosocial behaviors and traits (Donald et al., 2019): Multiple psychometric studies have shown that trait mindfulness is correlated with agreeableness (Thompson & Waltz, 2007), empathy (Dekeyser et al., 2008), and conscientiousness (Thompson & Waltz, 2007; Giluk, 2009); and mindfulness-based interventions and training programs are found to effectively enhance compassion and empathy (Kreplin et al., 2018; Campos et al., 2019).

Some prevailing frameworks for understanding mindfulness theorize that the practice improves attentional control, self-awareness, metacognition, and emotional control (Lutz et al., 2007; Vago & Silbersweig, 2012). Indeed, mindfulness-related changes in brain and behavior have been observed in attentional paradigms (Brefczynski-Lewis et al., 2007; Farb et al., 2007; Hasenkamp & Barsalou, 2012). and various socially relevant behavioral paradigms, such as the Affective Stroop Task (Allen et al., 2012), pain-related tasks (Grant et al., 2011; Lutz et al., 2013; Mascaro et al., 2013), and emotional provocation (Taylor et al., 2011). For example in one prior study, we tested prosocial decision-making using the Ultimatum Game as a function of mindfulness training (Kirk et al., 2016), and found that individuals who have gone through the training showed an increased willingness to cooperate. Taken together, these findings suggest that by training these cognitive functions, mindful individuals may be better able to observe and alter their social responses and emotional awareness, and develop prosocial characteristics (for a review, see Tang et al., 2015). Neuroimaging studies have further supported this claim of enhanced self-awareness and emotional control in mindfulness meditators: A meta-analysis found meditators, compared to novices, exhibited consistent changes in regions that have been associated with self-awareness (Craig, 2004), higher-order self-processing (Cavanna & Trimble, 2006), and metacognition (Christoff & Gabrieli, 2000; Fleming et al., 2010; McCaig et al., 2011). Electroencephalogram (EEG) studies, in turn, have identified lower frontal gamma activity in long-term meditators during task transition (Lutz et al., 2004).

Crucially, however, despite mindfulness’ association with prosocial traits and with various affective modalities in controlled laboratory studies, to our knowledge *no studies to date have examined the neural correlates of mindfulness in daily social interaction*. Here, we thus sought to investigate whether neural signatures of mindfulness identified in highly-controlled tasks extend into real-life social interactions, i.e. whether mindfulness-related traits predict neural responses during social interaction similar to those in laboratory-based tasks.

To assess trait mindfulness, we used the Mindful Attention Awareness Scale (MAAS), a standardized questionnaire designed to assess the open awareness of the present moment (Brown & Ryan, 2003). Mindfulness is theorized to have five distinct facets (Baer et al., 2006): observing (attending to internal experiences), describing (labeling inner experiences with words), acting with awareness (paying attention to ongoing activity), non-judging of inner experience (non-evaluative stance towards thoughts and feelings), and non-reactivity to inner experience (allowing feelings to come and go without getting caught up in them). It is important to note that the MAAS has recently been criticized for not measuring the acceptance component of mindfulness (Sauer et al., 2013) or non-judgmental awareness (Baer et al., 2006). However, despite this shortcoming, the MAAS has been widely applied and shown to successfully probe specific aspects of mindfulness, such as acting with awareness (Coffey & Hartman, 2008), perceived inattention (Van Dam et al., 2010), and burnout and engagement (Kotzé & Nel, 2016). In addition, more comprehensive alternatives such as the Five Facet Mindfulness Questionnaire (FFMQ), a 39-item scale that measures all five facets of mindfulness, were unsuitable due to their length: The MAAS, a 15-item scale, was more suitable given the time constraint of our experimental setups.

To record participants’ brain activity we used Muse, a 4-channel EEG headband commercialized as a neurofeedback tool for mindfulness-based stress reduction training (MBSR; Hashemi et al., 2016). As a neurofeedback tool, Muse and its accompanying app have been reported to be effective in reducing stress in breast-cancer patients (Millstine et al., 2019), and improving well-being and attention (Bhayee et al., 2016). However, mindfulness-related EEG research using the Muse headset has generated mixed results. For example, one study using the Muse observed a significant increase in beta and gamma frequencies in the post-meditation sessions compared to pre-meditation (Karydis et al., 2018). Another study using the “calm score” computed by the Muse app (a proxy for mindfulness), however, failed to observe “calm score” changes in participants after a 1-month meditation intervention. Additionally, the “calm score” did not reflect participants’ increased trait mindfulness, as measured by the MINDSENS scale (Acabchuk et al., 2020).

Portable, wireless EEG headsets are increasingly used to conduct social neuroscience research in naturalistic settings, and specifically in so-called hyperscanning studies—studies that simultaneously measure the brain activity of multiple people interacting with each other. Using a range of metrics to quantify inter-brain connectivity (Ayrolles et al., 2021), inter-brain coupling has been linked to a variety of factors during both verbal and non-verbal social tasks (Czeszumski et al., 2020). During social interactions outside of laboratory environments, our group has previously found that inter-brain coupling is linked to social closeness, personal distress, and shared social attention (Dikker et al., 2017, 2021).

The first aim of the present study was to investigate whether laboratory findings on the neural correlates of mindfulness replicate during naturalistic social interaction in an EEG device that has been explicitly associated with mindfulness meditation. Specifically, we asked if more mindful individuals exhibited enhanced EEG alpha (8–12 Hz) and theta (4–8 Hz) power during face-to-face social engagement (Takahashi et al., 2005).

The second aim of this research was to capture possible “multi-brain” neural correlates of mindfulness during naturalistic interaction. We investigated whether inter-brain coupling correlates with mindfulness during naturalistic social settings, building on a growing body of research on mindfulness on the one hand, and social neuroscience research using portable EEG systems on the other.

## Methods

### Participants

We included participants who partook in the Mutual Wave Machine exhibition at Espacio Telefónica in Madrid, Spain (2019; see Experimental Setup). 554 individuals participated in the study, including 271 females, 245 males, and three individuals who identified as “other”.

Participants’ ages ranged from 12 to 81 years, with an average of 33.8 years. Participants completed the questionnaires and consent forms in Spanish. As an art experience, participation was voluntary and without monetary compensation. Individual written informed consent disclosing the scientific purpose was obtained before the experimental session.

### Experimental Setup

This study was conducted as part of the participatory art installation Mutual Wave Machine, a Brain-Computer Interface (BCI) that translates real-time correlation of pairs’ EEG activity into light patterns (described in detail in Dikker et al., 2021; also see wp.nyu.edu/mutualwavemachine). This setup allowed us to study real-world face-to-face social interactions in a large population of participants recruited outside of the traditional research subject pool. Museum and gallery visitors freely interacted with each other while their EEG was recorded using the Muse, a four-electrode wireless EEG system (Krigolson et al., 2017). The interaction typically lasted ∼10 minutes. Real-time inter-brain power correlations were calculated and used to generate visual feedback for the participants as part of the 10-minute experience. Before the session, participants completed the MAAS, Inclusion of the Other in the Self Scale (IOS) Scale (see Materials), and the Personal Distress subscale of the Interpersonal Reactivity Index (IRI; see Materials). After completing the session, participants were seated back to the setup station and asked to fill out an additional set of questions, including the IOS Scale, IRI questionnaires, and the Positive and Negative Affect Schedule (PANAS-X; see Materials).

### Materials

All participants were asked to complete short questionnaires both before and after the session, addressing their relationship to each other, mood, and personality traits. Relationship measures include questions about relationship duration and social closeness. Affective personality trait measurement consisted of (a) a revised 14-item version of the Interpersonal Reactivity Index after the session (Davis & Others, 1980), including the subscales Personal Distress (e.g., “When I see someone who badly needs help in an emergency, I go to pieces”) and Empathic Concern (e.g., “I often have tender, concerned feelings for people less fortunate than me”); and (b) the MAAS (Brown & Ryan, 2003), consisting of 15 items measuring one’s awareness of what is taking place at the present (e.g. “I could be experiencing some emotion and not be conscious of it until some time later.”). Both questionnaires were answered on a five-point Likert scale ranging from “Does not describe me well” to “Describes me very well”. Social closeness was assessed using the Inclusion of the Other in the Self (IOS) Scale, a pictorial measure of closeness with two overlapping circles representing the self and the other (Aron et al., 1992). Participants also completed a shortened version of the Positive and Negative Affect Schedule (PANAS-X; Watson & Clark, 1994), which was not analyzed here.

### Data analysis

#### Personality Metrics

After removing incomplete and incorrect data entries, 475 individuals’ answers were preserved. For the purpose of this study, the following metrics were analyzed: MAAS score, social closeness scale, Personal Distress, Empathic Concern, sex, age. To investigate which trait measure was related to mindfulness, we constructed a multiple linear regression analysis using Personal Distress, Empathic Concern, age, and social closeness scale as predictors, and the MAAS score as the predicted variable.

#### EEG Preprocessing

The initial dataset consisted of 277 pairs of ∼10 minute recordings. First, EEG data files were removed if files were not readable (4 pairs), the two EEG files were misaligned (119 pairs), or subjects’ self-reported information was missing (57 pairs). Each individual EEG dataset was then bandpass filtered from 0.1 to 30 Hz, and segmented into 1-second epochs (“pseudo trials”). Bad channels were manually rejected upon visual inspection; artifacts in epochs were detected and rejected using the Python package “autoreject” (Jas et al., 2017). Finally, the automatic selection and correction were manually checked. This resulted in the additional exclusion of 20 pairs due to poor data quality. Note that in inter-subject connectivity analyses, we only preserved epochs that survive the preprocessing for both subjects in each pair: unmatched epochs were removed. For individuals’ power analyses, we used all the artefact-free epochs, regardless of the participants’ partners’ data, and removed participants with less than 60 clean epochs (60 seconds). Lastly, datasets with less than 50 remaining epochs (50 seconds) after these preprocessing steps were excluded from further analysis (15 pairs removed). These preprocessing steps resulted in 62 pairs (124 individuals) for the intersubject connectivity analysis, and 379 individuals for the power analysis.

After preprocessing, we performed the short-time Fourier transform on the 1-second epochs, using a Hanning window with a one-sample step size, resulting in complex spectral coefficients of 1 Hz resolution from 1 to 30 Hz.

#### Inter-brain coupling analysis

Inter-brain coupling was calculated using Circular Correlation Coefficient (CCorr), which is the phase synchrony measure that is argued to be robust to spurious synchronization (Burgess, 2013; Goldstein et al., 2018). CCorr was computed between corresponding channels in the dyad, and then averaged across channel pairs.

To capture slower, transient information throughout the time series, the epoched complex coefficients were concatenated before the correlation (henceforth referred to as “concatenated”), resulting in two discontinuous complex series from the pair. CCorr was then computed by correlating the angular component of the two concatenated series for each pair with respect to all four channels using the Python package Astropy (Astropy Collaboration et al., 2018). The computation is demonstrated in eq.1.1 (“Circular Correlation and Regression,” 2001), where *X* and *Y* are concatenated series from a certain frequency bin, and n represents the total number of time points (e.g. if 100 epochs are preserved, there are 256 Hz x 100 s = 25600 time points).

We also used a second metric, concatenated Projected Power Correlation (PPC) was calculated based on past studies (Dikker et al., 2021; Hipp et al., 2012) demonstrated in eq.1.2-5, where *X*(*t, f*) and *Y*(*t, f*) are the concatenated complex coefficients at frequency *f*. First, the projection of *Y*(*t, f*) on *X*(*t, f*) is removed, leaving only the part of *Y*that’s orthogonal to *X*, i.e. *Y*_⊥*X*_(*t, f*), and the same computation was done for *X*(*t, f*), resulting in *X*_⊥*Y*_(*t, f*) (eq.1.2). Second, we computed the correlation between |*X*(*t, f*)| and *Y*_⊥*X*_(*t, f*), and |*Y*(*t, f*)| and *X*_⊥*Y*_(*t, f*) respectively, and then averaged the two values as our PPC per frequency bin.

To investigate the difference between concatenating epochs versus averaging epochs, we also applied the same method to the epoched complex coefficients without concatenating (henceforth referred to as “epoched”). In such “epoched CCorr” or “epoched PPC”, the same calculation was done to individual epochs, and the result was an average of CCorr values across epochs. These exploratory results can be found in Figure S1 and Figure S2.

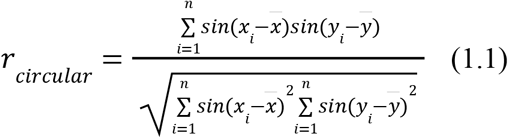

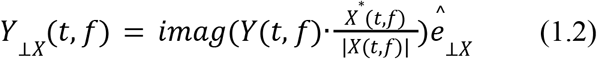

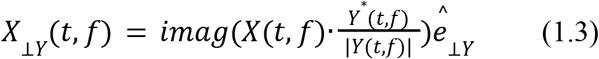

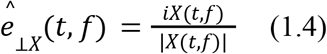

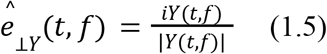

#### Cluster-based permutation analysis for correlation

To investigate the relationship between mindfulness and inter-brain coupling, we computed the Pearson Correlation Coefficient between pairs’ MAAS score and connectivity metrics in every frequency bin, using a cluster-based nonparametric test to correct for multiple comparisons. The protocol is adapted from Dikker et al. (2021). First, Pearson correlation coefficients were computed between every frequency bin and pairs’ mindfulness scales, generating 30 correlation values. Correlation significance thresholds r_upper and r_lower were then determined by choosing the 97.5th and 2.5th percentile of the 30 correlation values, respectively. Then the random permutation procedure started with randomly shuffling the behavioral variable (e.g. mindfulness scale), and computing correlation values for each frequency bin. Correlation values higher than r_upper or lower than r_lower were marked as significant in this permutation, and significant correlation values that were adjacent in frequencies were identified as clusters. For each cluster, we extracted the cluster size as our cluster statistics. When there was more than one cluster, the maximum cluster size was chosen. This random permutation procedure was repeated 2000 times, generating a distribution of cluster statistics, of which the 95th percentile was determined as the significant cluster threshold. Last, from the actual correlation values, we identified clusters using a p-value threshold of 0.1, and compared their sizes to the significant cluster threshold. The monte-carlo p-value was then calculated from the percentile score of the actual cluster size in the cluster statistic distribution. A monte-carlo p-value lower than the significant threshold 0.05 would mean the actual cluster size is larger than 95% of the random distribution of cluster sizes. We applied the same analysis to mindfulness’ correlation with both individuals’ EEG power and inter-brain coupling.

## Results

### Individuals’ EEG power does not predict mindfulness

Contrary to prior findings illuminating neurobiological changes related to mindfulness in laboratory contexts, i.e. a global increase in alpha and theta power during various kinds of meditations (Lee et al., 2018; Lomas et al., 2015; Takahashi et al., 2005), we found no significant correlation between individuals’ power spectral density during social interaction and their MAAS responses (Fig.1).

**Figure 1.**
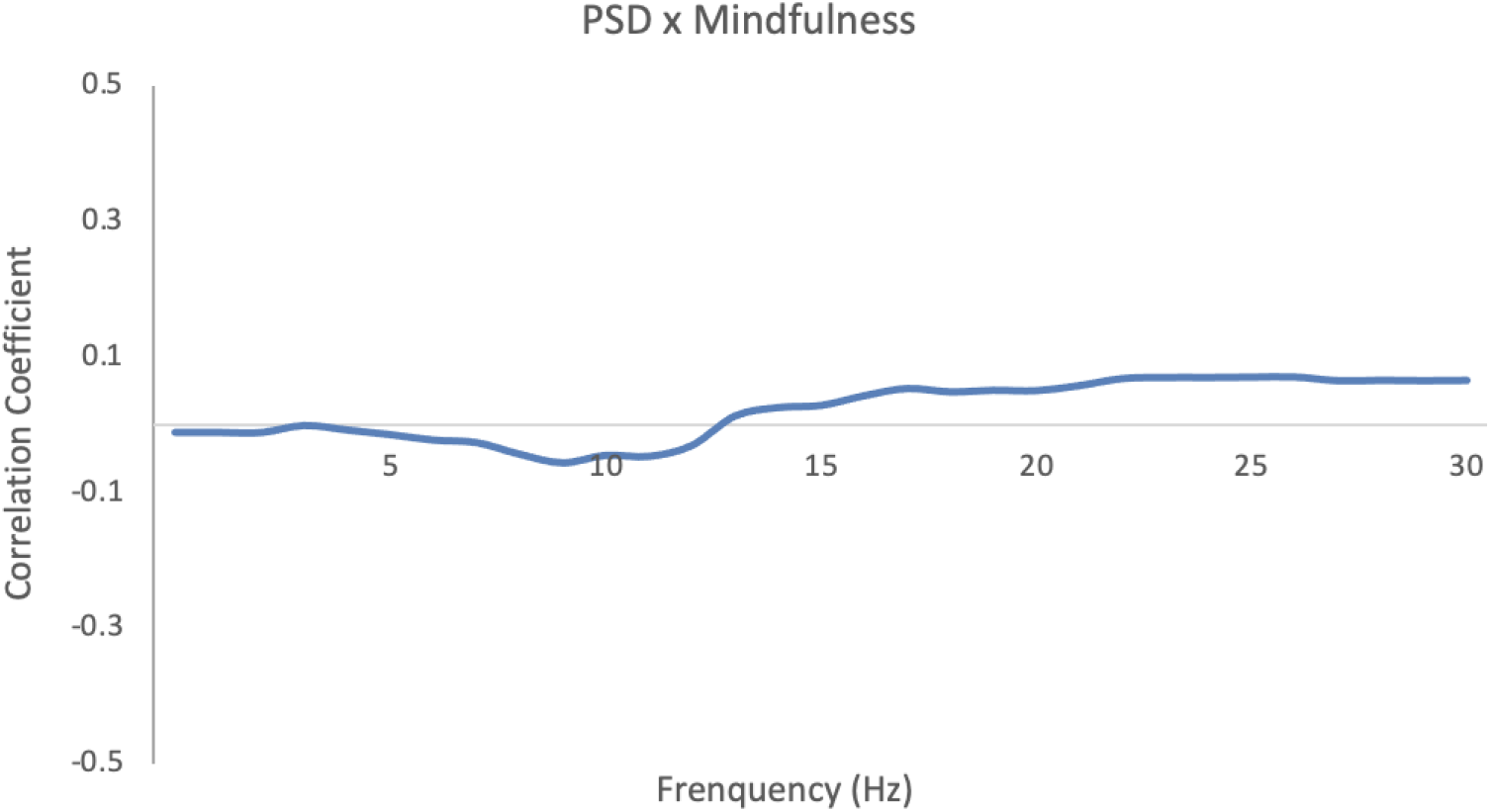
Pearson correlation coefficient between individuals’ MAAS score and their power spectral density (PSD). Cluster-based permutation analysis of the correlation coefficients showed no significant clusters (p=0.92).

### Inter-brain coupling is correlated with dyads’ mindfulness

The cluster-based permutation analysis showed that pairs’ mindfulness predicted inter-brain coupling (CCorr; monte-carlo p-value < 0.001). Specifically, as can be seen in Figure 2b, there were two clusters where the CCorr theta band (5-8 Hz) cluster shows a negative correlation between CCorr and subjects’ mindfulness (Fig.2b, circular correlation coefficient at 7 Hz; r(62) = - 0.373; p = 0.003), and the high beta band (26-27 Hz) cluster shows a positive correlation (Fig.2c, circular correlation coefficient at 26 Hz; r(62) = 0.325; p = 0.011). Figure 2a shows the Pearson Correlation Coefficient between the MAAS and CCorr at every frequency from 1 to 30 Hz, with significant frequency clusters marked with bold lines.

**Figure 2.**
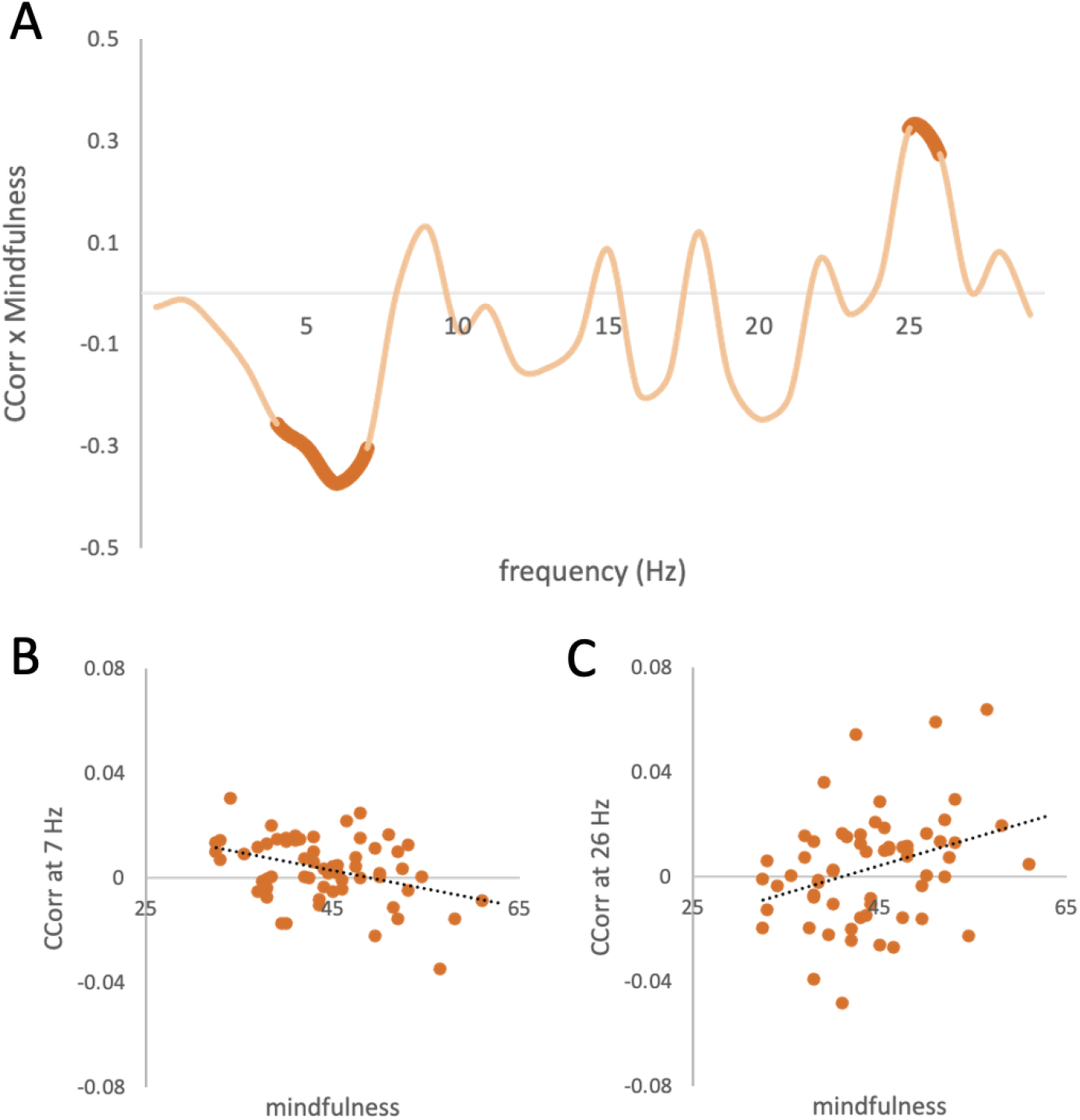
Correlation results between trait mindfulness and CCorr. (A) Pearson correlation coefficients (y-axis) between pair-averaged MAAS and inter-brain coupling (CCorr), for each 1-Hz frequency bin from 1-30 Hz (x-axis). Two significant clusters (p = 0.002) are highlighted in bold. (B) Scatter plot between CCorr at 7 Hz and pair-averaged MAAS. The dotted line is the linear regression line. (r(62) = -0.373; p = 0.003) (C) Scatter plot between concatenated CCorr at 26 Hz and pair-averaged mindfulness scale (r(62) = 0.325; p = 0.011)

### Personal Distress predicts mindfulness

The regression analysis showed Personal Distress as the only significant predictor (t(475) = -5.493, p < 0.001; Fig.3) for trait mindfulness (for the full results, please see Table S1). Individuals with lower personal distress reported higher mindfulness scale.

**Figure 3.**
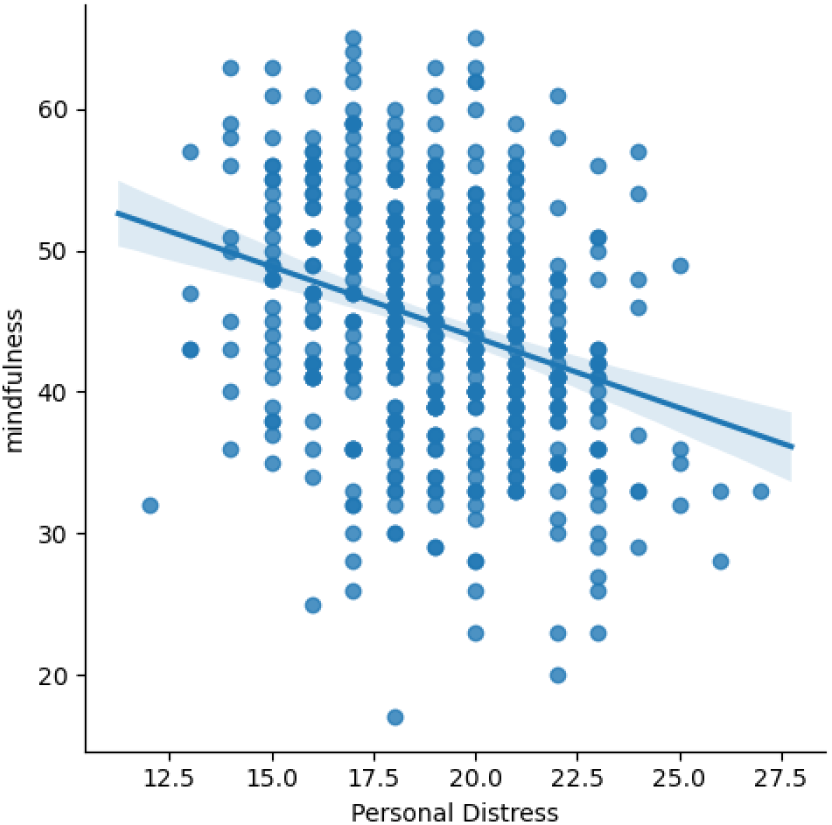
A linear regression model for personal distress and mindfulness. Individuals’ personal distress is negatively correlated with mindfulness scale.

## Discussion

This study investigates the neural correlates of mindfulness in daily social interactions. Our study investigated the relationship between mindfulness and prosociality on both the psychometric and neurobiological level. The questionnaire results showed that trait mindfulness was associated with lower personal distress. The EEG results showed that neural correlates of mindfulness during social interaction were found in intersubject synchrony (CCorr), but not in individual EEG power changes.

### Intra-vs inter-individual neural correlates of mindfulness

Contrary to previous studies, we did not find a relationship between individual brain activity (power spectral density) and mindfulness. There are a few possible explanations for this null effect. First, past studies investigating EEG power and mindfulness have been focusing on different types of mindfulness correlates from ours: power changes in individuals have been observed only in meditative states (Lutz et al., 2004; Takahashi et al., 2005), and functional changes in other controlled lab tasks are observed with fMRI (Allen et al., 2012; Hasenkamp & Barsalou, 2012; Lutz et al., 2013; Mascaro et al., 2013) or in event-related potentials with EEG (Brown et al., 2013; Wong et al., 2018). Second, to our knowledge no studies investigating the neural correlates of mindfulness have used EEG during dynamic social interaction. Finally, using a four-channel portable EEG system in a noisy, less controlled setting might also have contributed to a null result. It is important to reiterate, however, that we *did* find inter-brain correlates of mindfulness during social interaction.

This is not the first study to report a discrepancy between intra- and inter-brain neural correlates: other hyperscanning studies have similarly found that a multi-brain approach captures neural correlates of social behaviors that are not observed in individuals (Simony et al., 2016; Balconi et al., 2017; Davidesco et al., 2019). For example, in one study an inter-brain network but not individual brain activity predicted players’ strategy in prisoner’s dilemma (De Vico Fallani et al., 2010), and inter-brain coupling but not individual alpha power nor intra-brain synchrony predicted students’ performance during lessons (Davidesco et al., 2019). Under the rationale that online mutual interaction is “a complex nonlinear system that cannot be reduced to the summation of effects in single isolated brains” (Koike et al., 2015), our study further validates a multi-brain approach in complex social tasks by extending it to a naturalistic setting and portable EEG systems.

### Mindfulness and empathy measures

Our questionnaire results showed a negative correlation between trait mindfulness and personal distress, but not between trait mindfulness and empathic concern (see Results). In line with previous research, this result suggests that mindfulness might alleviate the negative consequences of empathy, but not necessarily contribute to empathic feelings: Previous research has suggested a complicated relationship between mindfulness and empathy, especially when empathy is dissected as a multidimensional construct (Davis & Others, 1980). Measuring overall self-reported empathy, some studies have reported a positive correlation between trait mindfulness and empathy (Beitel et al., 2005; Dekeyser et al., 2008; Greason & Cashwell, 2009), and mindfulness-based stress reduction (MBSR) training has been shown to increase participants’ self-reported empathy (Shapiro et al., 1998). However, other studies did not find effects of mindfulness-based training on self-reported empathy, empathic concern, or emotion recognition (Galantino et al., 2005; Lim et al., 2015). Specifically, in an eight-week MBSR training for nursing students, self-reported personal distress decreased, whereas self-reported empathic concern and perspective taking did not change (Beddoe & Murphy, 2004).

Distinct from empathic concern, personal distress is a self-oriented negative feeling that is often associated with reduced perspective taking and compassion fatigue when witnessing others’ suffering, and can be unrelated to prosocial behaviors (FeldmanHall et al., 2015; Pulos et al., 2004). Thus, our finding that trait mindfulness is negatively correlated with personal distress but not related to empathic concern, is in line with previous work. Such empirical evidence supports the theory that mindfulness is a vital part of self-compassion, which contributes to the resiliency against emotional fatigue (Figley, 1995; Neff, 2003; Thomas, 2012). Further evidence to bolster this argument comes from a study where participants displayed increased willingness to cooperate after having completed MBSR training, as indexed by higher acceptance rates to unfair monetary offers in the Ultimatum Game. Although no data was collected pertaining to participants’ personal distress, it is not unlikely that better distress regulation in the MBSR group when presented with unfair monetary offers in the Ultimatum Game may have contributed to this finding (Kirk et al., 2016).

### Relationship between trait mindfulness and inter-brain coupling

We observed a positive correlation between inter-brain coupling and mindfulness in the beta frequency range, but a *negative* correlation between trait mindfulness and inter-brain coupling in the theta frequency range. This finding seems puzzling in light of past studies reporting a positive relationship between inter-brain coupling and prosocial behavior and prosocial traits (cooperation, social closeness, joint attention, etc.; Valencia & Froese, 2020). However, recent work has pushed back on the leading assumption that more synchrony is always ‘better’ (for review, see Mayo & Gordon, 2020). For example, multiple studies have found cognitive downsides of behavioral synchrony, such as insecure attachment (Feniger-Schaal et al., 2016), worse performance in cooperative problem solving (Abney et al., 2015; Wallot et al., 2016), and decreased self-regulation (Galbusera et al., 2019). Physiological research has also yielded mixed results about the link between synchrony and couples’ relationships as well as parent-infant engagement (Timmons et al., 2015; Wass et al., 2019). In their review, Mayo and Gordon (2020) proposed situating synchrony in an interpersonal system that contains both collective and independent behaviors, and taking into account both the synchronization and segregation aspects inherent to synchrony. While neural studies overwhelmingly report positive relationships between inter-brain synchrony and social factors, there too, some studies pointed to the complexities of such relationships. For example, Goldstein et al. (2018) found that, during hand holding, romantic partners’ inter-brain coupling (CCorr) was *negatively* correlated with analgesia of the target person upon pain stimulation, and *positively* correlated with empathic accuracy of the partner. Separate examination of the effect of pain and touch, however, suggested distinct brain-coupling components associated with the experience of pain and the empathy for pain. The relationship between inter-brain synchrony and mindfulness, a personality trait that entails a mixture of cognitive properties, may also correspond to multiple processes and require further investigation.

## Conclusion

This study used consumer-grade portable EEG (Muse) and asked how trait mindfulness and prosociality manifest itself in neural responses during naturalistic dyadic face-to-face social interactions. Neural correlates of mindfulness during social interaction were evident in inter-brain coupling, but we failed to replicate previous studies showing individual EEG power changes as a function of mindfulness. In addition, the directionality and signature of the relationship between mindfulness and inter-brain coupling varied by frequency and by how inter-brain coupling was computed. Together, our findings are suggestive of a complex relationship between mindfulness and inter-brain coupling, while at the same time raising a cautionary note about methodological approaches in hyperscanning research.

