## Supplementary Information for "Trait mindfulness predicts inter-brain coupling during naturalistic face-to-face interactions"

| Source | B | SE B | t | p |
| --- | --- | --- | --- | --- |
| sex | 0.264 | 0.732 | -0.323 | 0.718 |
| (post) Personal Distress | -0.867 | 0.155 | -5.608 | <0.001 |
| (post) Empathic Concern | 0.191 | 0.137 | 1.390 | 0.165 |
| (avg) Social Closeness | 0.146 | 0.239 | 0.611 | 0.541 |
| age | 0.057 | 0.031 | 1.869 | 0.062 |

Table S1. Regression table for predicting individuals’ mindfulness scale. Both Personal Distress Scale and the Empathic Concern Scale were calculated from post-session answers to the Interpersonal Reactivity Index (IRI; Davis & Others, 1980); social closeness was an average of pre- and post-session answers to the Inclusion of the Other in the Self (IOS) Scale (Aron et al., 1992).

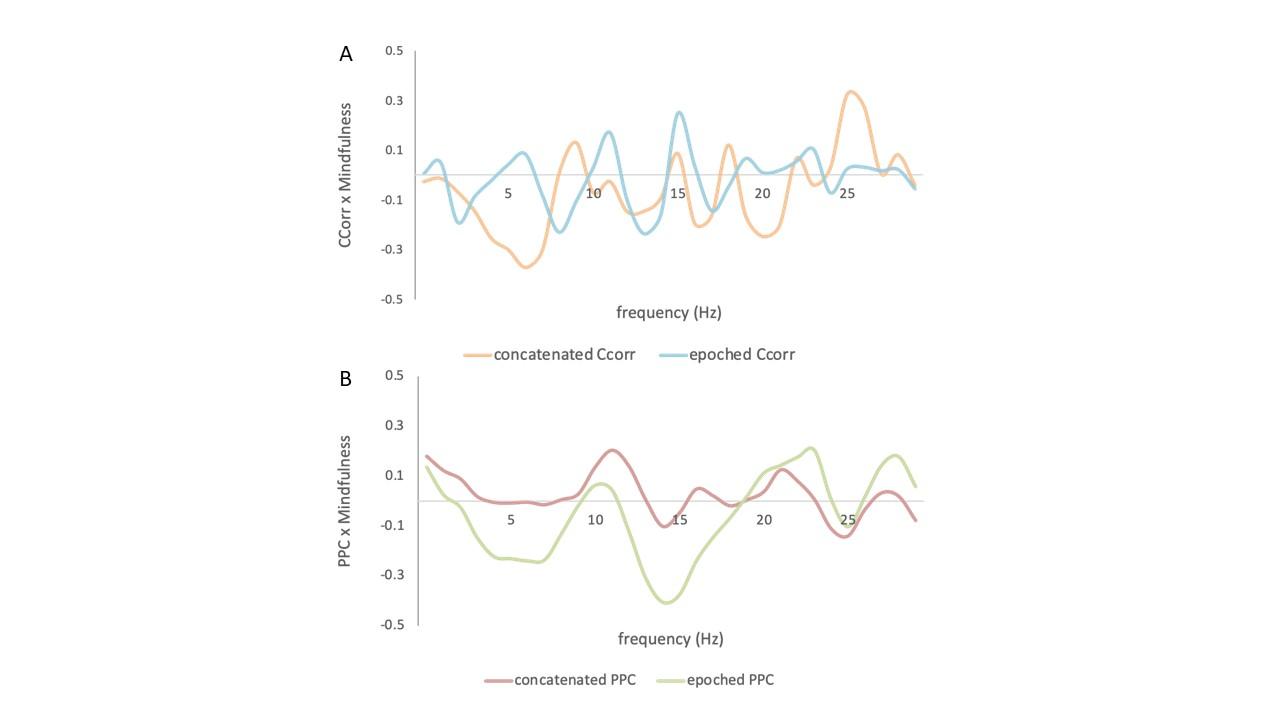

Figure S1. Pearson correlation coefficient between pair-averaged mindfulness and inter-brain synchrony, computed with (**A**) epoched Circular Correlation (CCorr; Burgess, 2013) and concatenated CCorr and (**B**) epoched CCorr and concatenated Projected Power Correlations (PPC; Hipp et al., 2012). While the averaging method does not affect PPC trend significantly, it does in the case of CCorr, showing different and even opposite relationships at certain frequencies.

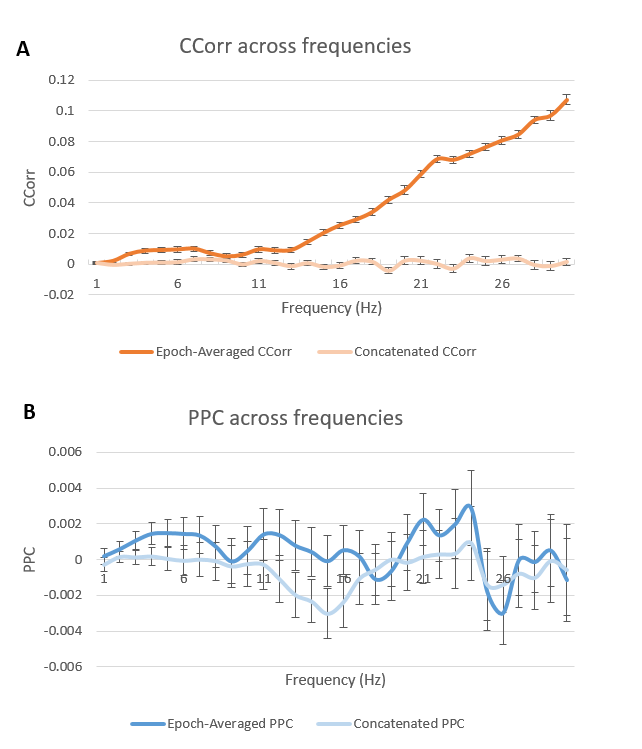

Figure S2. Connectivity values averaged across subject pairs for each frequency bin. (**A**) epoched and concatenated Circular Correlation (CCorr; Burgess, 2013). (**B**) epoched and concatenated CCorr.

As frequency increases, epoched CCorr values diverged significantly from concatenated CCorr, while epoched Projected Power Correlation (PPC; Hipp et al., 2012) and concatenated PPC stay relatively similar. This would explain the different correlation results in Figure S1, where epoched PPC and concatenated PPC had similar trends of correlation with mindfulness, whereas epoched CCorr showed different and sometimes opposite trends from concatenated CCorr. This interaction might be because phase synchrony captures much faster temporal information in epoched data compared to power synchrony as discussed in methods, and thus is more susceptible to window lengths. We thus suggest that phase synchrony computation should adopt frequency-dependent windowing or other methods to compensate for an increasing number of cycles as frequency rises in a fixed time period.
